# Rapid assessment of phytoplankton assemblages using Next Generation Sequencing – Barcode of Life database: a widely applicable toolkit to monitor biodiversity and harmful algal blooms (HABs)

**DOI:** 10.1101/2019.12.11.873034

**Authors:** Natalia V. Ivanova, L. Cynthia Watson, Jérôme Comte, Kyrylo Bessonov, Arusyak Abrahamyan, Timothy W. Davis, George S. Bullerjahn, Susan B. Watson

**Author notes:** Institut national de la recherche scientifique, Centre - Eau Terre Environnement, Québec, QC, Canada. National Microbiology Laboratory, Public Health Agency of Canada, Guelph, ON, Canada. Biology Department, University of Waterloo, Waterloo ON, Canada. These authors contributed equally to this work (Natalia V. Ivanova, L. Cynthia Watson). Corresponding author: (NVI).

## Abstract

Harmful algal blooms have important implications for the health, functioning and services of aquatic ecosystems. Our ability to detect and monitor these events is often challenged by the lack of rapid and cost-effective methods to identify bloom-forming organisms and their potential for toxin production, Here, we developed and applied a combination of DNA barcoding and Next Generation Sequencing (NGS) for the rapid assessment of phytoplankton community composition with focus on two important indicators of ecosystem health: toxigenic bloom-forming cyanobacteria and impaired planktonic biodiversity. To develop this molecular toolset for identification of cyanobacterial and algal species present in HABs (Harmful Algal Blooms), hereafter called HAB-ID, we optimized NGS protocols, applied a newly developed bioinformatics pipeline and constructed a BOLD (Barcode of Life Data System) 16S reference database from cultures of 203 cyanobacterial and algal strains representing 101 species with particular focus on bloom and toxin producing taxa. Using the new reference database of 16S rDNA sequences and constructed mock communities of mixed strains for protocol validation we developed new NGS primer set which can recover 16S from both cyanobacteria and eukaryotic algal chloroplasts. We also developed DNA extraction protocols for cultured algal strains and environmental samples, which match commercial kit performance and offer a cost-efficient solution for large scale ecological assessments of harmful blooms while giving benefits of reproducibility and increased accessibility. Our bioinformatics pipeline was designed to handle low taxonomic resolution for problematic genera of cyanobacteria such as the *Anabaena-Aphanizomenon-Dolichospermum* species complex, two clusters of *Anabaena* (I and II), *Planktothrix* and *Microcystis*. This newly developed HAB-ID toolset was further validated by applying it to assess cyanobacterial and algal composition in field samples from waterbodies with recurrent HABs events.

## Introduction

Outbreaks of harmful algal blooms (HABs) dominated by toxigenic and nuisance cyanobacteria are increasingly reported at the global scale [1–5] with adverse effects on the health, resilience of aquatic food-webs and many negative socioeconomic impacts, such as decreased water quality, recreation, businesses and property values [6–8]. Under high nutrient concentration dominance of cyanobacteria is associated with reduction of phytoplankton biomass resulting in lower zooplankton community diversity affecting aquatic food-webs [9–11]. HABs have garnered significant national and international attention, yet their management remains a major problem, as these events and their associated risks are difficult to identify and predict in a timely fashion. Expedient detection and accurate identification of toxigenic and bloom-forming species are essential to assess the potential risks associated with a bloom development, to identify the main sources of HABs taxa and to evaluate the main factors that drive their spatial and temporal dynamics. This information is fundamental to any effective management plan developed to predict, manage, and reduce HAB frequency, severity, and toxicity.

Traditionally, cyanobacteria and eukaryotic microalgae have been classified and identified by microscopic analysis of key morphological/cellular characteristics such as pigmentation, cell arrangement and size (unicell/filament/trichome/colony), specialised cells (heterocytes/akinetes/zygospores), gas vacuoles and sheath, cell wall, flagella, plastid number and arrangement, division planes etc. [12–15]. However, many of these diagnostic characters (e.g. size, colonial configuration, gas vacuoles, specialised cells) vary under different environmental conditions and can be lost during cultivation [16,17]. Komárek & Anagnostidis [15] note that up to 50% of strains in culture collections do not correspond to diagnostic characters of the taxa to which they were initially assigned. Because of these issues with traditional identification methods, cyanobacterial systematics have been undergoing widespread revision using a polyphasic approach, combining molecular analysis of 16S rRNA gene and other markers [18] with biochemical and other traits [19]. 16S rDNA is commonly used to identify algae and cyanobacteria and has been applied in DNA barcoding of harmful cyanobacteria [20], phylogenetic evaluation of heterocytous cyanobacteria [21], and accessing symbiotic cyanobacteria community in ascidians [22]. The DNA barcoding utilizes sequence diversity in short standardized gene regions for species identification and discovery [23] and although the use of DNA barcodes for species identification has been increasing, there are still no comprehensive reference libraries for freshwater phytoplankton, especially for toxin-producing species. As a fundamental part of this project outcome, we developed a curated reference database with focus on bloom-and toxin producers, generated for cyanobacterial 16S and algal 16S chloroplast rDNA. This database was derived from culture collections and hosted in Barcode of Life Data System (BOLD) [24], an analytical workbench and depository for DNA barcodes linking voucher specimen information (collection data and digital images) with sequence data, including laboratory audit trail and sequence trace files.

Given the socioeconomic importance of HABs, a rapid method for community-wide phytoplankton assessment offers an important tool to detect and monitor for bloom-forming and toxigenic taxa and serve as an effective early warning system for the development of potentially harmful blooms. Molecular techniques such as quantitative PCR (qPCR) have been used for the rapid detection of toxigenic and bloom-forming cyanobacterial and algal species [25–28], but this technique is limited to a few species at a time. NGS offers an alternative and potentially more powerful metagenomic approach to rapidly and accurately identify multiple species from a mixed sample. This approach has been used successfully in previous studies to assess environmental samples for eubacterial, cyanobacterial and phytoplankton composition [28,29] and diatom species assemblages [30], and for evaluating methodological biases in mock communities [31–33]. Yet, all studies to date have been conducted alone with their own sets of primers and experimental conditions, moreover, there is still no standardized and comprehensive database of cyanobacterial and phytoplankton sequences which is urgently needed.

Here we report the results of a multi-year study designed to develop a HAB-ID toolset for rapid assessment algal and cyanobacterial diversity with focus on two important indicators of ecosystem health: toxigenic bloom-forming cyanobacteria and impaired planktonic biodiversity using a combination of DNA barcoding and NGS. This work systematically addressed four main objectives (Fig 1): 1) construction of a reference BOLD database for algal and cyanobacterial 16S ribosomal RNA, using isolates from the Great Lakes and other regions; 2) optimization of Polymerase Chain Reaction (PCR) primers and DNA extraction protocols; 3) validation of the NGS method on mock communities generated from known cyanobacterial and algal cultures; 4) application of the HAB-ID tool to environmental samples and comparison of its performance with traditional microscopy.

**Fig 1.**
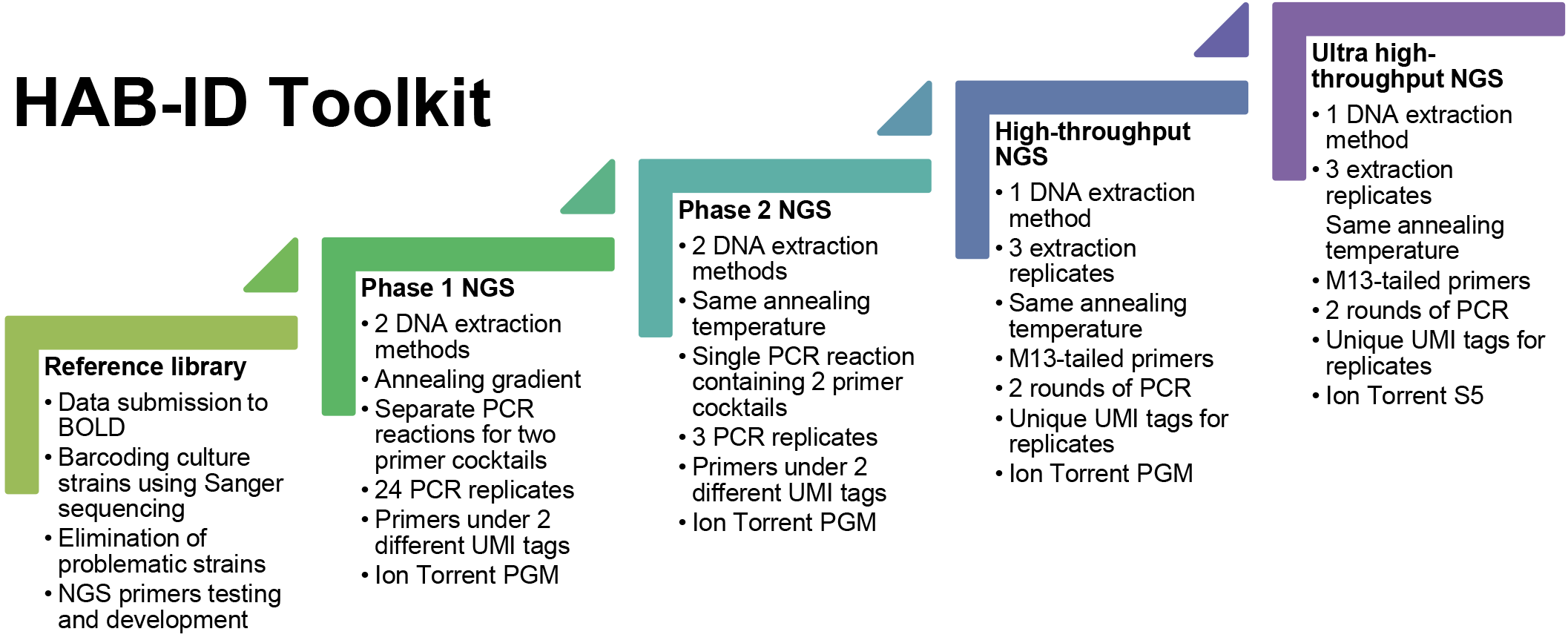
Project phases - from initial optimization to high-throughput analysis of environmental samples

## Material and methods

### Ethics statement

Environmental samples were collected from permanent Environment and Climate Change Canada (ECCC) survey stations in Lakes Erie, St. Clair and Winnipeg. No specific permissions were required for the sampling stations/activities because the sampling stations were not privately owned or protected. This study did not involve endangered or protected species.

### Strains and culture conditions

The 203 of 263 analyzed cyanobacteria and algal strains used in this study are listed in public BOLD dataset: DS-ECCSTAGE (https://doi.org/10.5883/DS-ECCSTAGE) along with collection information and microscopic images. Strains which failed to sequence of did not pass validation were moved into private problematic samples project ECPS (Table S1).

Of these strains, 91 were collected from the Laurentian Great Lakes and other waterbodies throughout Canada and first isolated into sterile filtered lake water using sterile micropipetting and repeated washings. Once established they were transferred to growth media. and housed at the Canadian Centre for Inland Waters in Burlington, Ontario (CCIW). Cyanobacteria stock cultures were maintained as batch cultures in 55 ml tubes at 18°C in Z8 media [34] at a 16:8 h light:dark light regime of 50 μmol photons m^-2^ s^-1^. Eukaryotic strains were maintained at 14 ± 1°C in Chu10 [35], WC [36] or BBM [37] at a 16:8 h light:dark light regime of 110 μmol photons m^-2^ s^-1^. All strains were maintained as monoalgal (non-axenic) batch cultures and transferred bimonthly or monthly using sterile techniques. To augment the database, additional strains were obtained from the following culture collections: the Canadian Phycological Culture Collection (CPCC; Waterloo, Canada), the Metropolitan Water District of Southern California (G. Izaguirre), the University of Zurich (F. Juttner), the Norwegian Institute for Water Research (NIVA; Oslo, Norway), the Pasteur Culture Collection (PCC; Paris, France), the Polar Cyanobacteria Culture Collection (PCCC; Quebec, Canada), the Culture Collection of Algae at Göttingen University (SAG; Göttingen, Germany), and the University of Texas Culture Collection (UTEX; Texas, USA).

All strains for DNA barcoding were pelleted and stored in Lysing Matrix A tubes containing 1 ml of ddH_2_O at −80°C until extraction.

### Preparation of mock communities

Mixes of 4-16 strains of cyanobacteria and algae were prepared in a single flask to produce mock communities using equal volumes from each culture regardless of density. A total of 9 mock communities were created in 2016 and 2017, which were split between two experiments: Mock1 and Mock2 (S1 Table). Subsamples of 20-40 ml from each culture mix were filtered onto 0.22 μm polycarbonate or polyethersulfone membranes in replicates, initially frozen at −80°C, transported on dry ice and kept at - 20°C prior to extraction. For some mock communities one subsample was preserved in Lugols’s solution to measure the relative proportion (by cell number) of each species using microscopy. To evaluate possible bias from DNA extraction and PCR amplification, mock communities for Mock community 1 experiment were extracted and amplified separately, then pooled for the first NGS run while for the second run DNA was pooled prior to PCR amplification. For second experiment (Mock community 2) each of the filters was split in half and extracted using 2 DNA extraction methods.

### Environmental samples – collection sites

Environmental samples were collected from three waterbodies which exhibit annual HABs due to excessive nutrient loading. Lake Erie, the smallest of the Laurentian Great Lakes, is of enormous economic value [8], providing drinking water for an estimated 11.5 million Canadian and American residents (IJC, 2014) and generating more than 7 billion dollars in revenue each year from fishing and tourism [8]. In the last couple of decades Lake Erie has been experiencing annual blooms of toxic cyanobacteria with increasing toxicity and duration [2,38,39]. Other areas of these Lakes (e.g. Lake St Clair) are also exhibiting frequent blooms, while the past decade or so has seen an alarming rise in HABs in Lake Winnipeg, dubbed the ‘sixth Great Lake’.

Samples were collected from four permanent Environment and Climate Change Canada (ECCC) survey stations in Lake Erie (879, 880, 885, and 970) in August 2015 and May 2016, five stations in Lake St. Clair (008, 134, 136, 139, and 142) in September 2016, and four stations in Lake Winnipeg (W1, W2, W5, and W8) in September 2017 (S2 Table). Water samples were collected from a depth of 1 m in Lakes Erie and Winnipeg and 0.5 m in Lake St. Clair using Niskin or van Dorn bottles. Water samples were filtered through Millipore 0.22 μm Sterivex filters (800-1000 ml) using a peristaltic pump, 0.22 μm polycarbonate (250 ml), or polyethersulfone (300 ml) membrane filters and immediately stored at −80°C for DNA extraction as outlined below. Subsamples from Lake Erie stations (879, 880, 885, and 970) collected in August 2015 were also processed for phytoplankton community composition and biomass (1% v/v final conc. Lugol’s iodine solution) for comparison purposes between NGS and microscopic identification.

### Microscopic analysis

Morphological identification and enumeration of phytoplankton in environmental samples were analyzed at the Algal Ecology & Taxonomy Inc. (ATEI; Winnipeg, Manitoba) with an exception for samples from station LE881 and LE885 collected on August 18, 2015, which were analyzed at Université du Québec à Montréal (UQAM) (Quebec). Cells were enumerated on a Leitz Diavert inverted microscope at 125-625X magnification (ATEI; Utermohl technique following [40] and an Olympus IMT-2 inverted microscope at 100-400X magnification (UQAM; Utermohl technique following [41], after sedimentation of a known volume of sample in a counting chamber. Taxa were identified to the lowest taxonomic level possible (usually species). For comparison with NGS, all identifications were used at the genus level, considering NGS naming convention used for species complexes and taxa with low resolution.

### Processing of samples for Sanger and NGS analyses

The processing procedures of environmental samples, cultures and mock communities are summarized in Table 1. Details about the different extraction protocols are given below.

**Table 1.**
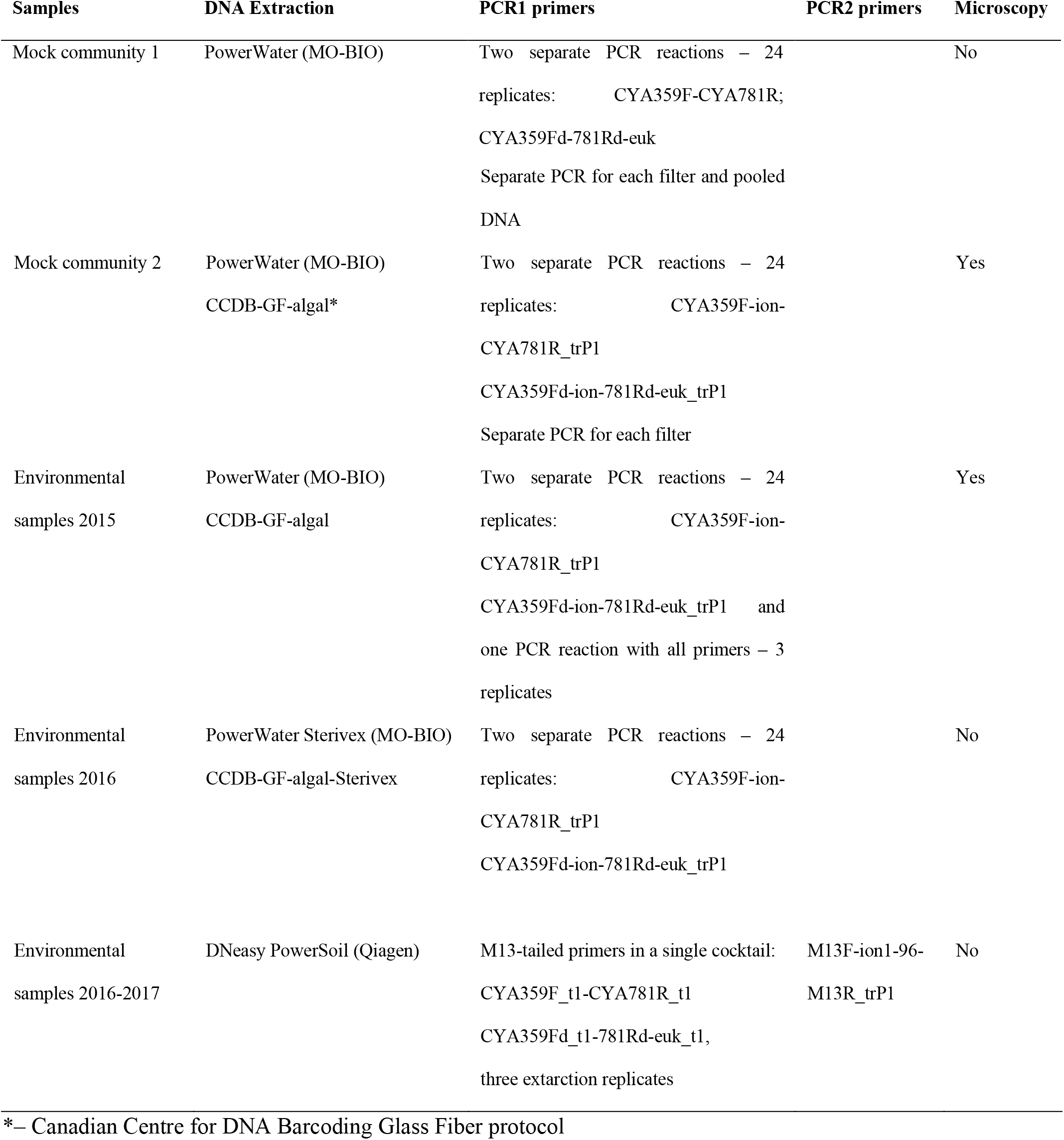
Summary of tested DNA extraction and PCR protocols for mock communities and environmental samples (detailed listing of strains included in the mock communities given in S1 Table).CCDB DNA extraction from algal cultures (CCDB-GF-cultures)

A volume of 50 μl of Proteinase K (20 mg/ml) and 200 μl of 5M GuSCN Plant Binding buffer were added to each tube containing pelleted algal culture in 1 ml of water. Tubes were briefly vortexed and incubated for 1 hour at 56°C on an orbital shaker. Tissue was further homogenized at 28 Hz for 1 min using TissueLyser (Qiagen) followed by 45 min incubation at 65°C. Tubes were centrifuged at 10,000×g for 2 min to pellet debris and 100 μl from each lysate was transferred to 1 ml Ultident tube rack; total genomic DNA was extracted as described in [42,43] with two WB washes; DNA was eluted in 65 μl of pre-warmed 10 mM Tris-HCl pH 8.0.

### CCDB DNA extraction from mock communities and environmental samples (CCDB-GF-algal)

Filters were placed in PowerWater grinding tubes; 50% bleach and ethanol, followed by flame sterilization were used to decontaminate instruments between samples. DNA was extracted as described in “CCDB-GF-cultures” with minor modifications. A volume of 50 μl of Proteinase K (20 mg/ml) and 1 ml of ILB buffer [42] were added to each tube, which were then briefly vortexed and incubated for 1 hour at 56°C on an orbital shaker. Tissue was further homogenized on Genie2 Vortex for 5 min at maximum speed followed by 45 min incubation at 65°C. Tubes were centrifuged at 4,000×g for 2 min to pellet debris and 150 μl from each lysate was transferred to 1.5 ml Eppendorf tube containing 300 μl of 5M GuSCN Plant Binding Buffer, and an entire volume was transferred to Econospin column (Epoch Life Science) for binding DNA to the membrane. DNA was extracted as described in [42] with two WB washes; DNA was eluted in 120 μl of pre-warmed 10 mM Tris-HCl pH 8.0. DNA was quantified using Qubit. DNA from mock communities was normalized to concentration 4 ng/μl prior to PCR; DNA from environmental samples did not exceed 4 ng/ μl and was used without normalization.

### Sterivex CCDB GF protocol for DNA extraction from environmental samples (CCDB-GF-algal-Sterivex)

A volume of 0.5 ml of ILB buffer and 50 μl of Proteinase K (20 mg/ml) were added to each Sterivex filter. Capped Sterivex filters were shaken for 5 min at minimum speed on Genie 2 vortex and incubated at 56°C for 1 hour. A volume of 1 ml of 5M GuSCN buffer was added to each tube, tubes were incubated at 65°C for 30 min and shaken 5 min at minimum speed. Tube contents were transferred with syringe to a new 2 ml tube; two loads of 400 μl were applied to Econospin column (Epoch Life Science) for binding DNA to the membrane (two loads had to be used due to volume capacity of 650 μl). The remainder of the procedures followed the methods described above under “CCDB-GF-Algal”. DNA was eluted in 100 μl of pre-warmed 10 mM Tris-HCl pH 8.0 and quantified using Qubit; DNA from environmental samples did not exceed 4 ng/ul and was used without normalization.

### PowerWater (MO-BIO) protocol for DNA extraction from mock communities and environmental samples

Filters were placed into PowerWater grinding tubes; 50% bleach and ethanol followed by flame sterilization was used to decontaminate instruments between samples. DNA extraction procedure was carried out in UV sterilized laminar hood using PowerWater DNA extraction kit according to manufacturing instructions using alternate lysis protocol with incubation at 65°C for 15 min. DNA was eluted in 100 μl of PW6 buffer, quantified using Qubit and normalized to concentration 4 ng/μl prior to PCR.

### Sterivex PowerWater (MO-BIO) and DNeasy PowerSoil (QIAGEN) protocols for DNA extraction from environmental samples

DNA extraction of Lake Erie samples was carried out in UV sterilized laminar hood using alternate lysis protocol with incubation at 65°C for 15 min for Sterivex PowerWater kits. Transfer volumes were reduced to 625 μl to minimize cross-contamination between samples. DNA was eluted in 100 μl of PW6 buffer and quantified using Qubit. Samples from Lakes Winnipeg and St. Clair were extracted with DNeasy PowerSoil kits (QIAGEN) at the ECCC following manufacturer’s protocol. This kit features standard 2 ml grinding media tubes, which fit into mini-centrifuge, and efficient removal of inhibitors removal, enabling DNA extraction from water and sediment samples. Therefore, we chose this kit for processing of field samples at the ECCC. Seventy percent ethanol and Eliminase followed by flame sterilization was used to decontaminate instruments between samples.

### Sanger sequencing workflow for reference library generation

The target genetic marker (16S) was amplified using PCR primers PLG1-1 and PLG2-1 to amplify ~1100-1200 bp of 16S and CYA106F-GC and CYA781R(ab) (Table 2) to amplify ~800 bp of 16S; followed by cycle sequencing with a standardized commercially available BigDye Terminator v3.1 kit to produce bidirectional sequences with partial sequence overlap. Sequencing reactions were analyzed by high-voltage capillary electrophoresis on an automated ABI 3730xL DNA Analyzer. DNA sequences recovered from the algal strains were deposited in project STAGE and corresponding public dataset DS-ECCSTAGE in the Barcode of Life Data System (BOLD) accessible at http://www.boldsystems.org/. After data validation, all samples found to represent contaminated or misidentified strains were moved into ECPS project on BOLD. After submission of eukaryotic strains with high sequence divergence BOLD systems algorithms were not able to handle data alignments and taxon ID tree generation, therefore data was processed in Geneious software v 11.0.4. Sequences were aligned using MAFFT v 7.308 [44,45], with the following parameters: Algorithm: Auto; Scoring matrix: 200 PAM/2; Offset value: 0.123. The Neighbor-joining tree was generated using Tamura-Nei distance model, exported as newick format and visualized in iTOL [46].

**Table 2.**
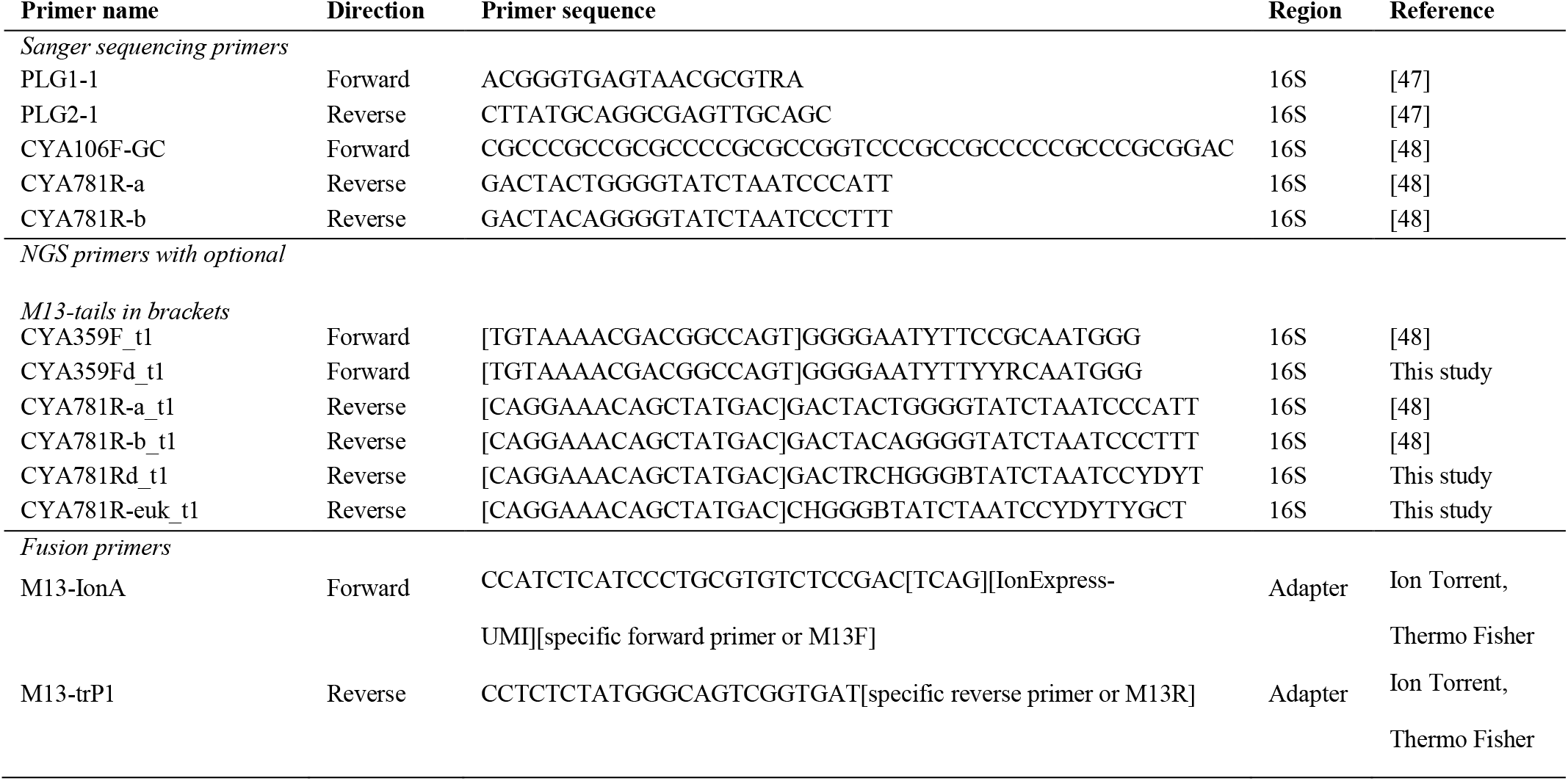
Primers used in Sanger and NGS workflows

### The Next Generation Sequencing (NGS) workflow

The target genetic marker 16S cyanobacterial rDNA or 16S chloroplast for eukaryotic strains was amplified using PCR for 30 cycles with fusion primers targeting shorter internal fragment of ~ 400 bp (Table 2) labelled with IonExpress UMI (Unique Molecular Identifier) tags using Platinum Taq as described in [49].

For the 24-replicate setup with annealing temperature gradient 2 μl of DNA from each sample was added to each 12 wells of the row into 96-well plates containing PCR mix with fusion primes with IonXpress UMI tags (Plate 1 – CYA359F -CYA781R, Plate 2 – CYA359Fd-CYA781Rd-euk), resulting in 24 PCR replicates per sample with each combination of primer pair and sample labelled with different UMI tag (after pooling each sample was represented by 2 UMI-tags). For the 3-replicate setup without annealing temperature gradient, each DNA sample was added in 3 replicates to a well with PCR master mix containing all primers (CYA359F-781R, CYA359Fd-CYA781Rd-euk), after pooling each sample was represented by 2 UMI-tags.

Single round PCR with fusion primers thermocycling consisted of: initial denaturation at 94°C for 2 min followed by 30 cycles (35 cycles for environmental samples) of: denaturation at 94°C - 1 min; annealing gradient (56-61°C) or annealing at 58°C for simplified PCR setup - 1 min, extension at 72°C for 1 min; final extension at 72°C for 5 min.

In the later high-throughput phase of the project, utilizing already optimized primer cocktails, samples from Lakes St. Clair and Winnipeg were extracted in three replicates using DNeasy PowerSoil kits and amplified with M13-tailed complete primer cocktail using two rounds of PCR. PCR1 products were diluted 2x and an aliquot of 2 μl was transferred to each well of PCR2 plate containing unique IonXpress UMI-tag to allow comparison of sample replicates.

PCR1 with M13-tailed primers thermocycling consisted of: initial denaturation at 94°C for 2 min followed by 20 cycles of: denaturation at 94°C - 1 min; annealing at 58°C- 1 min, extension at 72°C for 1 min; final extension at 72°C for 5 min.

PCR2 with M13 IonXpress1-96 primers thermocycling consisted of: initial denaturation at 94°C for 2 min followed by 20 cycles of: denaturation at 94°C - 1 min; annealing at 51°C- 1 min, extension at 72°C for 1 min; final extension at 72°C for 5 min.

Pooled amplicon libraries were purified using paramagnetic beads prepared as described in [50] using bead to product ratio 0.8:1. Each library was normalized to 13 pM for templating reaction. Ion PGM Template OT2 Hi-Q View kit, 316 or 318 chips and Ion PGM Sequencing Hi-Q View Kit for Ion Torrent PGM were used for sequencing, according to manufacturer’s instructions.

Only high-quality reads (QV>20 and longer than 200 bp) assigned to correct Ion Express MID tags were used in NGS data analysis. The following bioinformatics workflow was used to process NGS data: Cutadapt (v1.8.1) was used to trim primer sequences; Sickle (v1.33) was used for filtering (less than 200 bp were discarded) and Uclust (v1.2.22q) was used to cluster OTUs with 99% identity and minimum read depth of 10. Local BLAST 2.2.29+ algorithm was utilized to match resulting OTUs to custom reference library database for 16S generated from available BOLD records DS-ECCSTAGE dataset in BOLD; BLAST output results were parsed using custom built python script [51] OTUBLASTParser.py (available at https://gitlab.com/biodiversity/OTUExtractorFromBLASToutput): with the following 16S Identity Filter (Species 99%, Genus 97%, Family 95%, Order 90%); matches were concatenated using custom script ConcatenatorResults.py (available at https://gitlab.com/biodiversity/OTUExtractorFromBLASToutput), exported to Excel, and further processed in Tableau software 10.2 (min coverage 10).

NGS data for mock communities and environmental samples were submitted to ENA archive https://www.ebi.ac.uk/ena/browser/view/PRJEB27854.

### 16S data parsing to improve BLAST algorithm performance

Correct taxonomic assignments remain problematic for some cyanobacteria, especially for *Anabaena, Aphanizomenon, Dolichospermum* complex [52]. Moreover, *Microcystis* and *Planktothrix* have low interspecific divergence with 16S. Therefore, to ensure accurate identification with less false-positive species-level top hits using BLAST algorithm, we assigned sequences belonging to these groups to superficial taxonomic groups, based on Taxon ID tree (Fig 1). The annotated taxonomy file is available as Table S3.

## Results

### Reference library

All Sanger sequences for 16S rDNA were uploaded to BOLD, followed by validation of data using BOLD Taxon ID trees or neighbour-joining trees generated in Genious software. Strains with questionable placement were sent to experts for re-identification, re-ordered from culture collections or removed from the dataset. Our final validated BOLD reference database DS-ECCSTAGE contains a total of 203 strains of cyanobacteria and algae isolated from the Great Lakes and other areas, including nuisance and toxic species (Fig 2). Following sequence validation, sixty strains, most of which were contaminated or misidentified were transferred to private project on BOLD containing problematic samples (ECPS).

**Fig 2.**
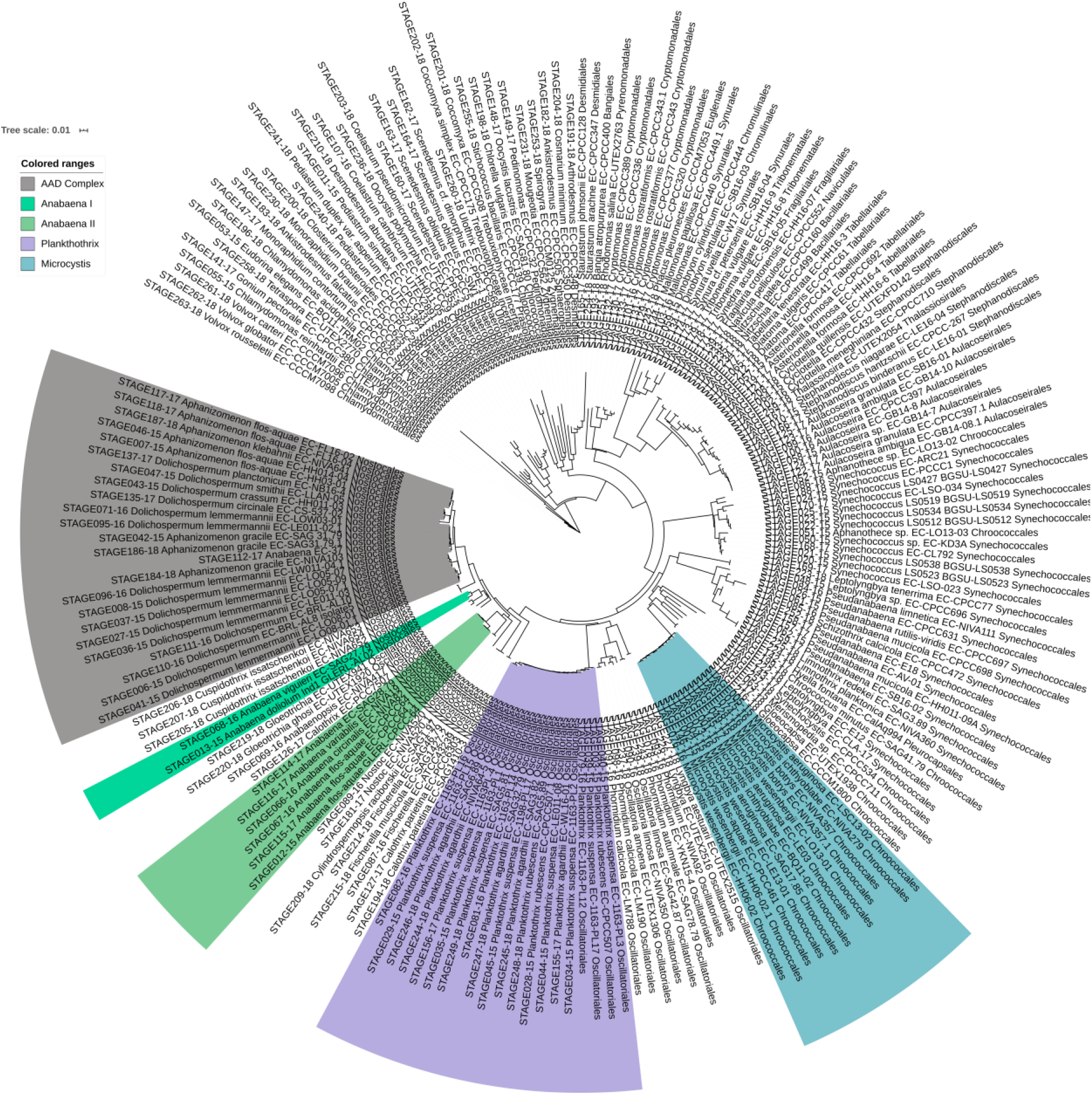
Neighbor-joining tree visualized in iTOL, showing taxa complexes and genera with low resolution used in BLAST search assignments and OTUparser.

As previously reported by [21], heterocytous cyanobacteria form a monophyletic group according to 16S rDNA sequence data; within this group, *Anabaena* and *Aphanizomenon* are intermixed. All planktonic *Anabaena* with gas vesicles were recently transferred into the genus *Dolichospermum* [52]. Such mixed groups represent a real challenge for accurate taxonomic assignments using BLAST-search algorithm. Therefore, we created groups, corresponding to closely related taxa (some with identical or nearly identical sequences), based on 16S data. In our study 24 strains of one *Anabaena,* five *Aphanizomenon* and fifteen *Dolichospermum* isolates formed an intermixed cluster which we named *Anabaena-Aphanizomenon-Dolichospermum* complex, hereafter called AAD Complex (Fig 2). From the *Anabaena* strains which did not cluster with the above-mentioned species complex, two (*A. doliolum* and *A. vigiueri*) were named *Anabaena I* and remaining strains, including *A. circinalis*, *A. variabilis* and *A. flos-aquae*, were named *Anabaena II*.

This validated reference database can be used as a local BLAST search database in NGS studies or for identification of strains sequenced using Sanger sequencing.

### Validation using mock communities

Mock communities assembled from cultures used to generate reference library (Table S1) enabled the validation of extraction protocols and minimization of PCR bias prior to working on field samples.

To evaluate primer bias and DNA pooling effect two NGS runs were completed on DNA isolated from mock community 1. For the first run, DNA from each filter was amplified separately and then pooled for the NGS run; for the second run - DNA was pooled prior to PCR setup. Our data demonstrated 99-100% efficient recovery of cyanobacteria and eukaryotic algae submitted for analysis, excluding problematic strains, which were removed from the reference database (Fig 3). All negative DNA extraction controls did not produce PCR products in any of mock community experiments.

**Fig 3.**
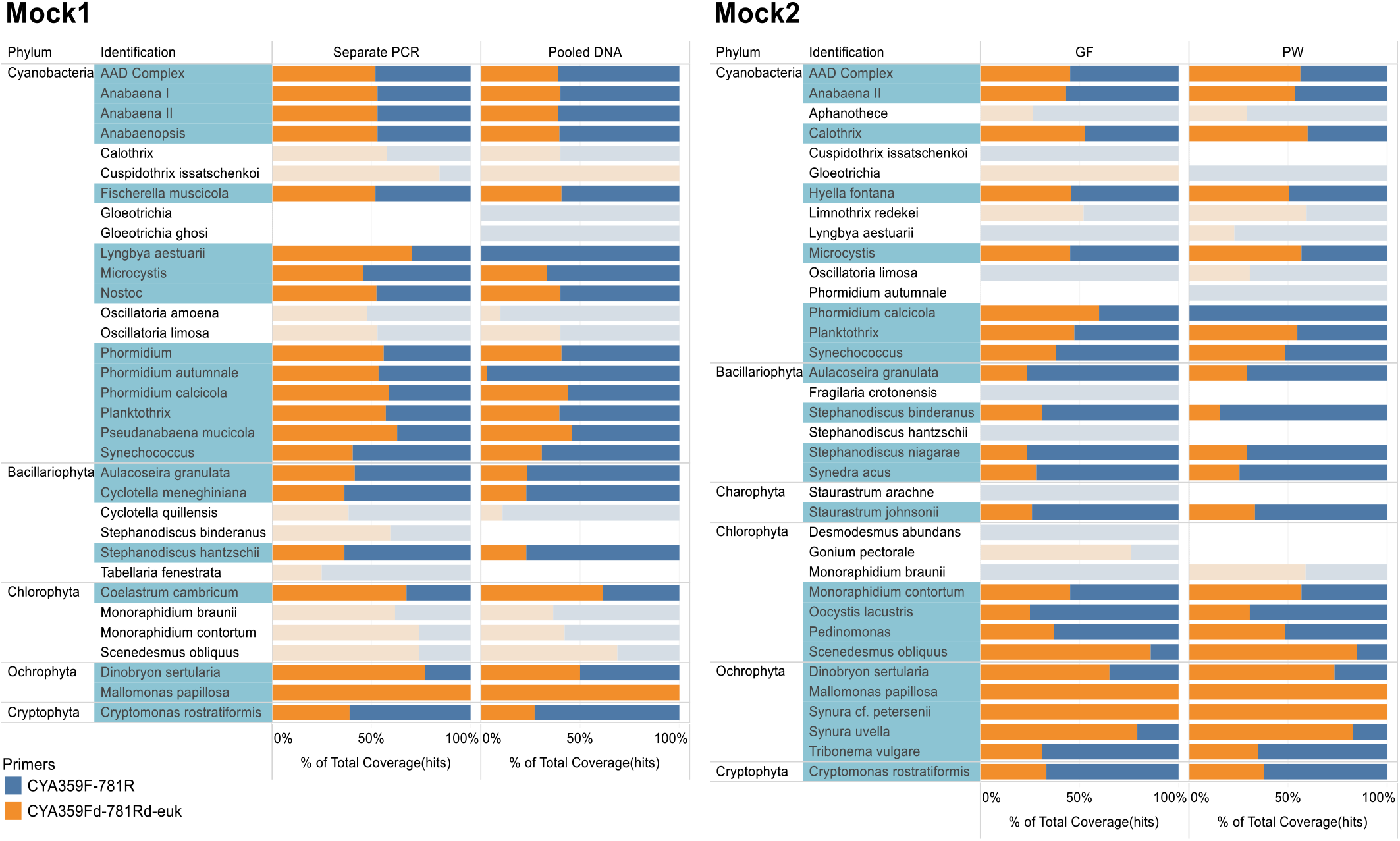
Primer specificity (Mock 1 and 2), number of PCR replicates (Mock1) and two extraction methods (Mock2) tested on mock communities. Shaded in blue – expected taxa; non-shaded – contaminant taxa.

While working on the first mock community dataset and reference library strains it was observed that some of the strains produced heterogeneous sequence data, resulting in background noise or failures in Sanger sequencing, or additional strains detected in mock community samples (Fig 3). Microscopy data (Fig 4) for 4 filters from mock community 2 confirmed that some of the analyzed strains contained contaminants of other cyanobacteria or algae, easily detected with NGS sequencing due to high sensitivity of the method. All problematic strains were removed from reference database and placed into private BOLD project ECPS.

**Fig 4.**
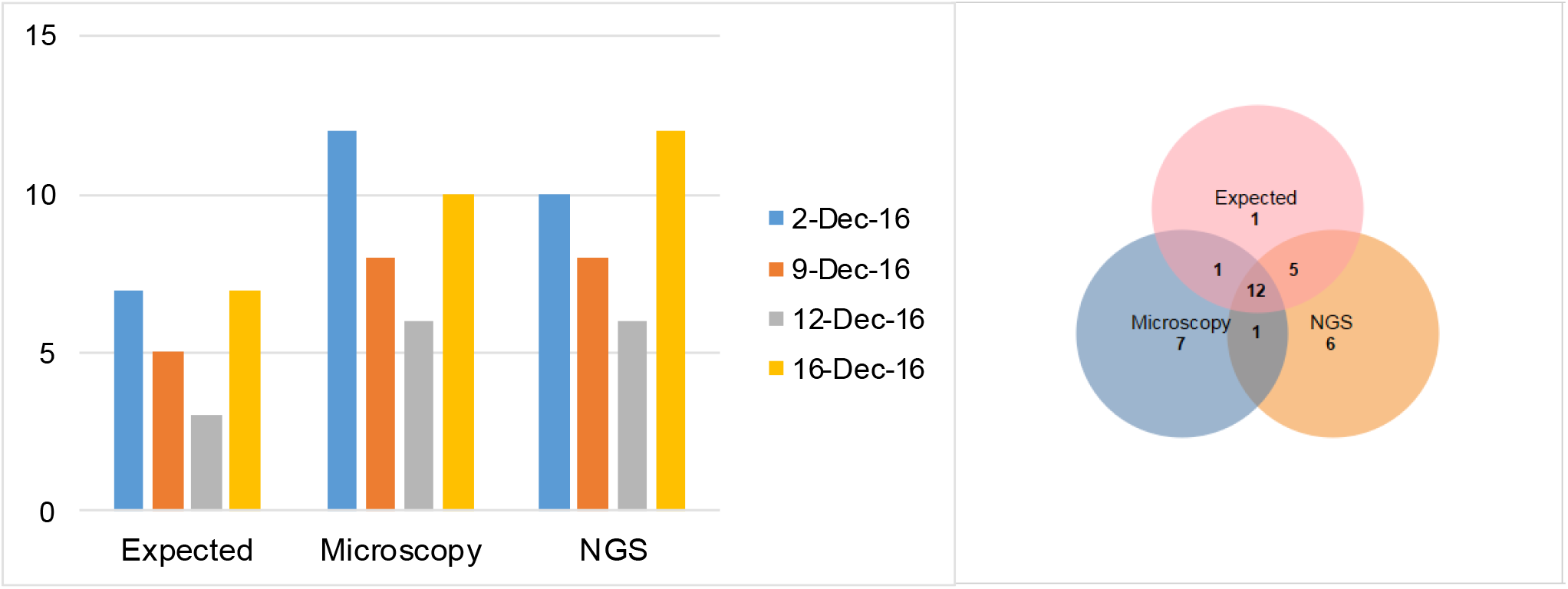
Genera count for 4 filters from mock community 2 (expected versus microscopy and NGS) and Venn diagram representing overall genera count of submitted strains from 4 filters and their detection using microscopy and NGS.

For Mock 1 experiment *Lyngbya* failed to be detected in pooled DNA, while *Phormidium autumnale* had less coverage. We tested two protocols for DNA extraction on mock community 2 and both protocols resulted in similar taxonomic recovery (Fig 3) with minor differences: *Phormidium calcicola* was not detected in PowerWater DNA extraction with CYA359Fd-781Rd-euk primer pair, while *Mallomonas papillosa* and *Synura cf. petersenii* were detected only using new modified primers CYA359Fd-781Rd-euk targeting eukaryotic algae in both DNA extraction protocols. *Euglena gracilis* from Mock community 2-Dec-16 (Table S3) failed to amplify both in Sanger sequencing and NGS workflows.

### Testing on environmental samples

The protocols were tested on environmental samples and resulted in recovery of 24 orders from environmental samples, where newly designed primer cocktail, targeting eukaryotic algae, improved recovery of Chlorophyta, Ochrophyta, Rhodophyta and Euglenozoa (Fig 5).

**Fig 5.**
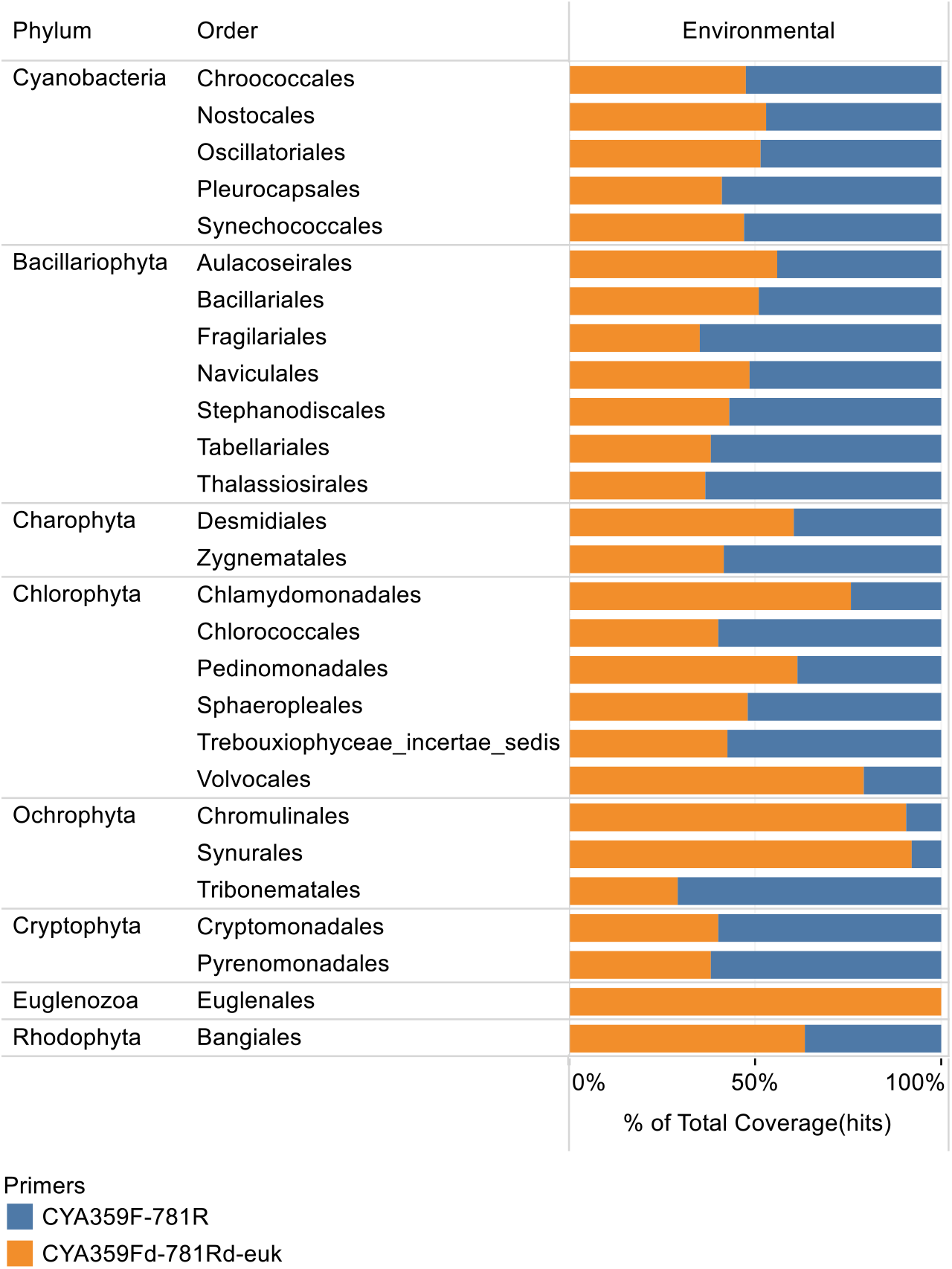
Nine environmental samples (n=4 – polycarbonate filters, n=5 - Sterivex filters) processed using two primer cocktails and two DNA extraction methods: overall order level recovery for two primer pairs (including order, family, genus and species level IDs)

Environmental samples from Lake Erie stations collected in duplicates on polycarbonate filters in August 2015 and on Sterivex filters in May 2016 were used for evaluation of two DNA extraction methods. Samples collected on polycarbonate filters were also used for evaluation of simplified PCR setup (with 3 PCR replicates) without annealing temperature gradient.

Overall, we were able to obtain comparable identification counts for two extraction treatments (Fig 6) and two PCR setups, as well as comparable read coverage profiles, except for 2 samples collected from station LE880 in 2016 submitted on Sterivex filters, which had different read coverage profiles between DNA extraction methods.

**Fig 6.**
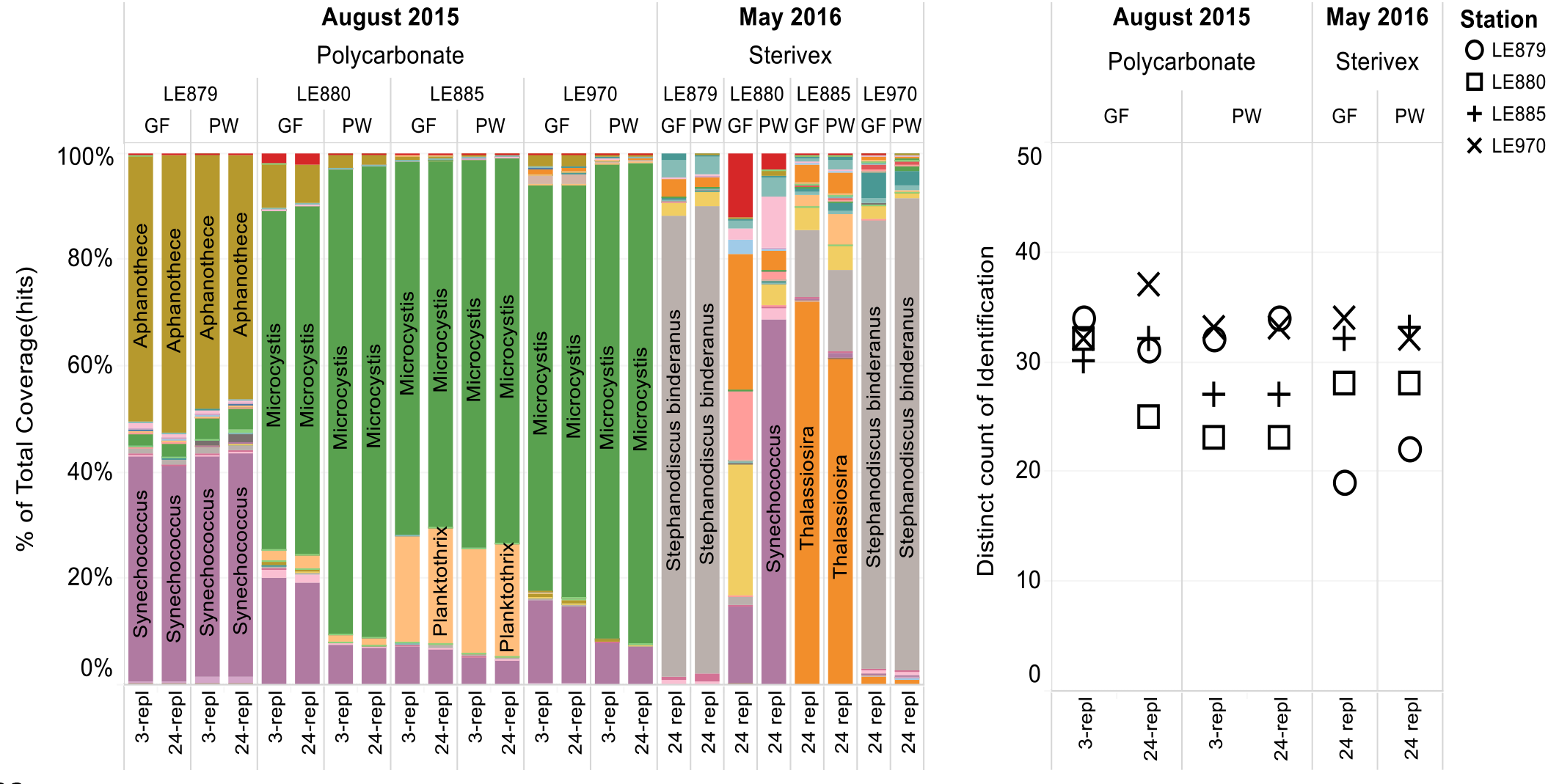
Read coverage (left) and taxonomic recovery for genus and species level identifications using NGS (right) from samples collected at four stations in Lake Erie tested using two DNA extraction methods. Samples collected from polycarbonate filters were also tested in two PCR protocols (3 and 24 PCR replicates).

The M13-tailed primers allow for the simplification and standardization of the workflow, while preserving leftovers of DNA for additional analyses. Furthermore, to accommodate for efficient multiplexing in a high throughput setting we added M13-tails to existing primers and combined all of them into a single primer cocktail in our current workflow, which was used for processing of environmental samples from Lake St. Clair and Lake Winnipeg. These environmental samples were extracted at ECCC using DNeasy PowerSoil kit and submitted for high-throughput analysis in 96-well plate format. UMI-tags were introduced using robotic PCR setup. While we continue to expand the reference library, our current database was successfully used to identify most of the reads present in environmental samples from Lake St. Clair and Lake Winnipeg (Fig 7).

**Fig 7.**
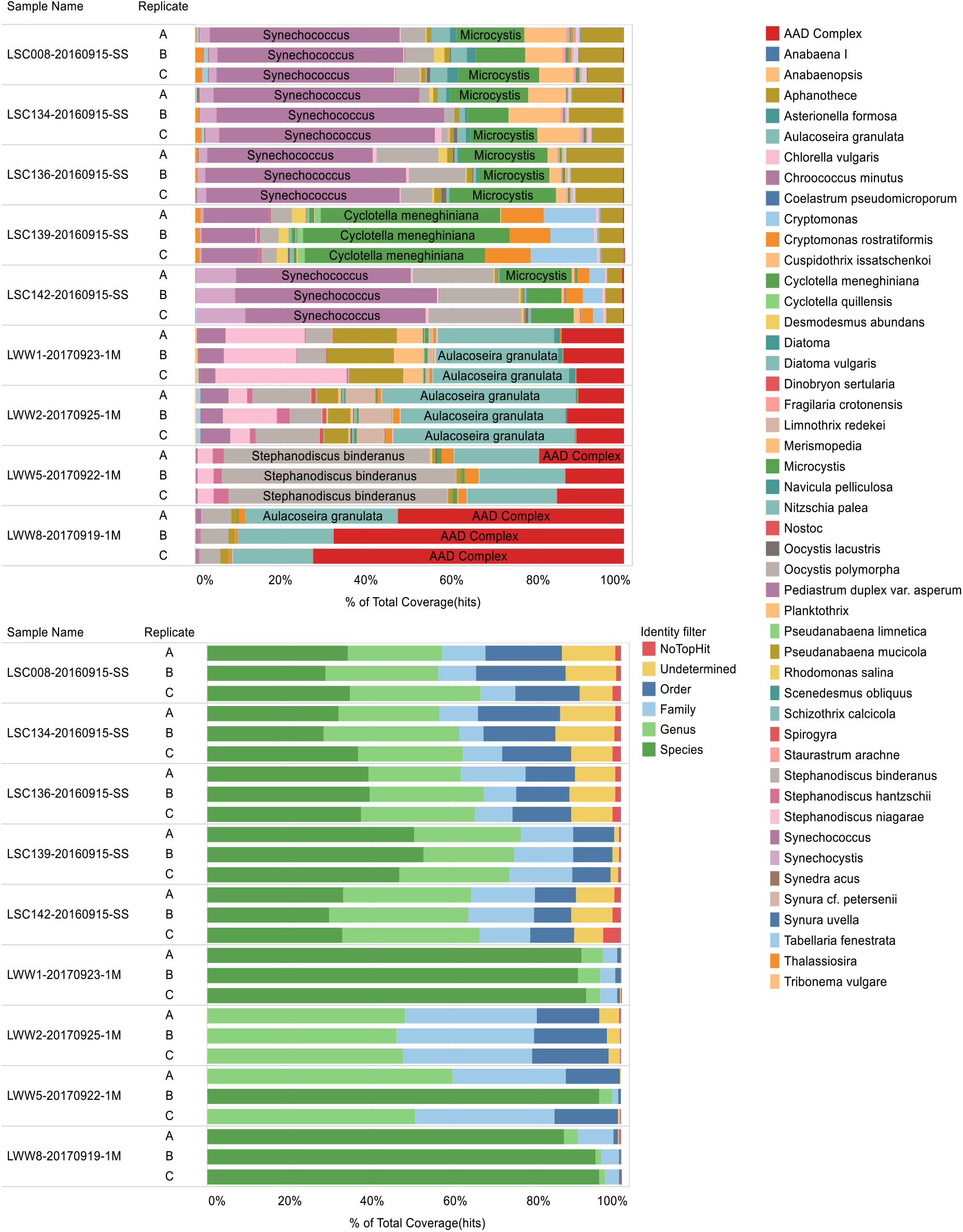
Environmental samples from Lake St. Clair and Lake Winnipeg collected and extracted in 3 replicates and amplified using M13-tailed primer cocktail. Top – read coverage for species and genus level identifications; bottom – distribution of identification top hits.

### Microscopy

Our reference database did not contain taxa from some phyla which were detected by microscopy. Therefore, we included only those phyla present in our database for direct comparison with NGS. The total number of genera for the four Lake Erie sites in August 2015 detected with microscopy and NGS are presented in Table 3. Overall, NGS detected significantly more genera than microscopy; across all samples, the number of detected genera ranged from 16 to 20 for microscopy and 30 to 36 for NGS.

**Table 3.** Number of cyanobacterial and algal genera for phyla present in the database detected using microscopy and NGS (only genus/species level ID counts) at four Lake Erie sites collected in August 2015.

|  | LE879 |  | LE880 |  | LE885 |  | LE970 |  |
| --- | --- | --- | --- | --- | --- | --- | --- | --- |
|  | Microscopy | NGS | Microscopy | NGS | Microscopy | NGS | Microscopy | NGS |
| August 2015 | 19 | 36 | 20 | 29 | 16 | 30 | 20 | 32 |

Cyanobacteria and Bacillariophyta genera were dominant in most samples followed by representatives of Chlorophyta (Fig 8). For all stations NGS detected more genera. According to both microscopy and NGS results Cyanobacteria were dominant in all samples collected in August 2015 during the bloom period.

**Fig 8.**
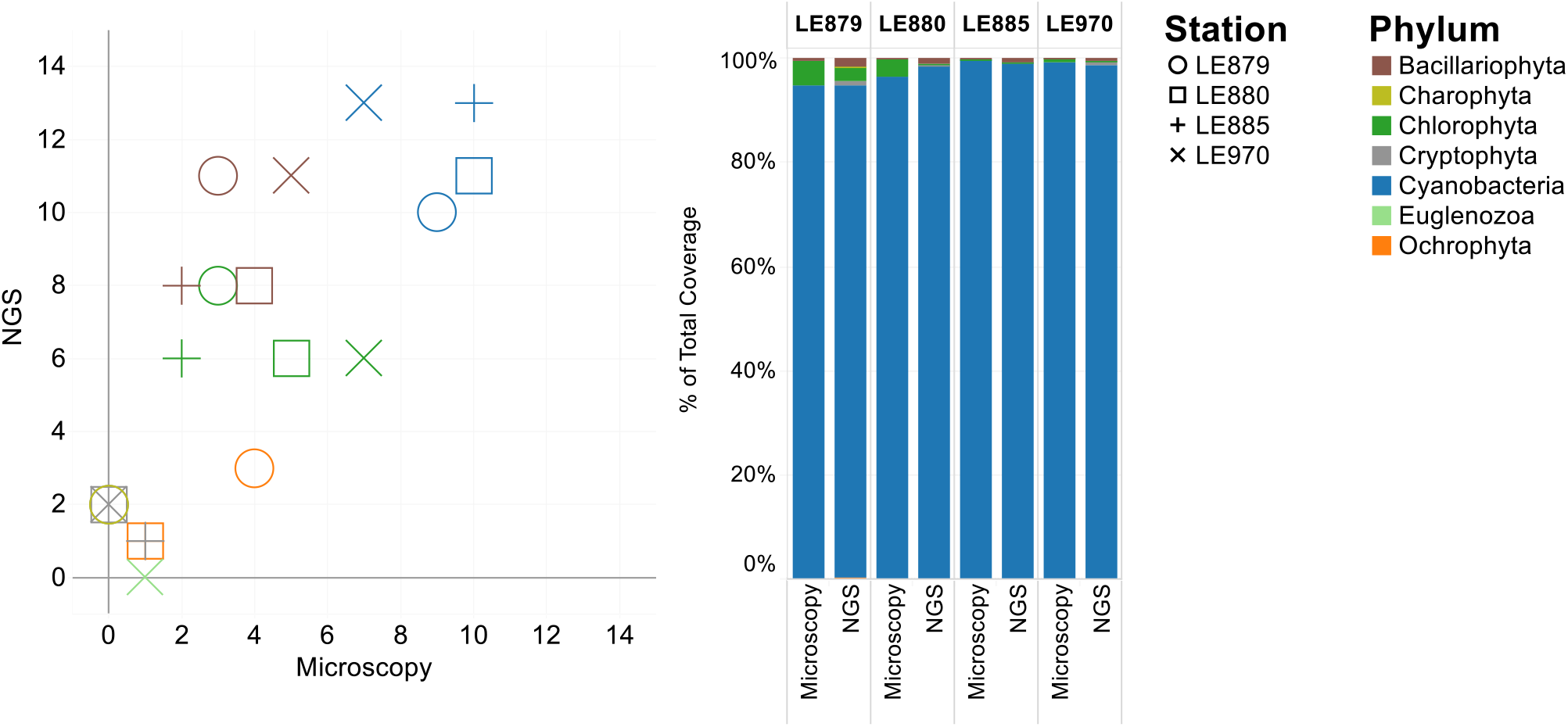
Comparison of genera counts detected by microscopy and NGS colored by phylum for samples collected in August 2015 at four stations in Lake Erie (left) and cell counts/NGS read coverage (right). Only phyla present in dataset were included in comparison.

## Discussion

In this work, we developed a HAB-ID toolset to detect toxigenic bloom-forming cyanobacteria and access community composition and diversity using a combination of DNA barcoding and NGS. Built around a diverse reference BOLD database dataset DS-ECCSTAGE for algal chloroplast and cyanobacterial 16S ribosomal RNA, the HAB-ID toolset successfully recovered distinct species in mock communities and provided accurate and more detailed taxonomic composition of natural phytoplankton in comparison to microscopic enumeration.

Isolating and maintaining monospecific strains of cyanobacteria and eukaryotic algae is not a trivial task, particularly with large culture collections. Strains can be mislabelled, lose morphological or biochemical traits or harbour contaminants present at isolation (e.g for cyanobacteria with significant sheaths), with implications for experimental work - as a result, axenic strains are very difficult to obtain [53]. According to the study of Cornet et al. [54] on contamination level of publicly accessible cyanobacterial genomes, 21 out of 440 surveyed genomes were highly contaminated (mostly with Proteobacteria and Bacteroidetes). In our study we used primers preferentially amplifying cyanobacterial 16S [47,48], however, some strains failed to sequence due to overlapping heterogeneous signal in Sanger sequencing, which can be indicative of contamination. Moreover, both microscopy and NGS detected algal/cyanobacterial contaminants in some of the cultures used to create reference library (Fig 4), and placement in the NJ tree also revealed that some of the strains had been wrongly identified by the culture collection. All such questionable strains and those, which failed to produce sequence data were removed from our validated reference dataset and placed into unpublished private BOLD project ECPS. The DNA from these strains can be accessed later using new technological advances such SMRT (Single Molecule Real Time) Sequencing using PacBio Sequel instruments, which allow accurate long reads via CCS (Circular Consensus Sequences) and, unlike Sanger sequencing enable separation of mixed signal in contaminated strains. These technological advances will enable to failure-track our DNA collection and recover more 16S rDNA sequences for BOLD reference database.

Some of traditional markers ntcA and rbcL [55,56] used for working with phytoplankton were less scalable for high-throughput processing due to lower amplification/sequencing success (data not shown). The 16S rDNA is commonly used for cyanobacteria [20,21,31,57] and, therefore, was selected as a primary marker due to its scalability for high-throughput screening. We chose internal 16S V3-V4 region for NGS analysis using existing primers for 16S of cyanobacteria [48] because this region contains significant information for phylogenetic assignments [58]. Furthermore, the length of the region is approximately 400 bp, which makes it compatible with Ion Torrent NGS sequencing platforms. Many of the previous studies on eukaryotic algae have utilized 18S rDNA instead of 16S [59–62], however, we chose 16S chloroplast rDNA as a single marker to enable simultaneous detection of cyanobacteria and eukaryotic algae for assay simplicity. While these primers are widely used in cyanobacterial studies [57,63,64], Zwart and co-authors [65] made modifications to these primers to increase their specificity. In our case, the availability of longer sequences in the reference database enabled us to verify primer binding sites, focus on strains underrepresented in NGS runs, and improve their recovery by introducing degeneracies in existing CYA359F – CYA781R primers or slightly shifting CYA781R primer position to enable detection of 16S chloroplast of eukaryotic strains. The most noticeable improvement of read coverage for eukaryotic strains in mock communities (Fig 3) and in environmental samples (Fig 5) using new primer cocktail CYA359Fd-781Rd-euk was detected for the following eukaryotic orders: Chlamydomonadales (75.82%), Volvocales (79.26%), Chromulinales (90.76%), Synurales (92.03%) and Euglenales (100%).

While commercially available Qiagen PowerWater and PowerSoil kits (currently available as QIAGEN DNeasy PowerWater and DNeasy PowerSoil) allow easy standardization between different labs, those protocols are time consuming. If processed in the centrifuge, PowerWater columns were difficult to transfer into clean tubes due to splashing of buffers and required frequent change of gloves. Because volumes listed in the protocol are close to the lid and can cause potential tube leaks and cross-contamination, we reduced the transfer volume to 625 μl. In this study, we detected contamination in only one negative extraction control extracted with PW kit. All contaminants detected in negative controls were filtered from the results for environmental samples. The CCDB GF extraction protocols utilize spin columns for binding high-quality DNA (Epoch Life Science) and custom buffers which allow efficient lysis of cyanobacteria and algae, and easy removal of inhibitors with reduced number of handling steps (i.e. the inhibitor removal step is omitted and only single load of sample is required for DNA binding). Contrary to MO-BIO PowerWater kit, these columns are easy to transfer into clean tubes and do not require frequent change of gloves. The protocol is scalable and available in 96-well format.

Overall CCDB extraction protocols (CCDB-GF-algal and CCDB-GF-algal-Sterivex) resulted in similar taxonomic recovery from mock community and environmental samples in comparison with MO-BIO PowerWater kit (e.g, total number of taxa detected using CCDB-GF extraction protocols and 24 PCR replicates for polycarbonate filters from 4 stations was 48 taxa versus 43 taxa for polycarbonate filters, and 37 vs. 40 with MO-BIO PowerWater kit for Sterivex filters (see also Figs 3, 5). The CCDB extraction protocols are therefore a good alternative to commercial kits that may be subject of manufacturing changes. In January 2016 MO-BIO was merged with Qiagen and we noticed better quality binding columns in DNeasy PowerWater and DNeasy PowerSoil kits. However, DNeasy PowerWater kit was reported to be incompatible with filters preserved in ethanol [66], which can be potential issue for remote sampling without immediate access to cold storage.

After evaluating primer specificity on mock communities and environmental samples we eliminated temperature gradient for annealing stage and reduced number of PCR replicates from 24 to 3, without noticeable impact on number of genera recovered and read coverage (Fig 1 and 6). The next optimization stage included incorporation of M13-tailed primers for high-throughput workflow, compatible with robotic plate-based PCR setup for incorporation of IonXpress UMI-tags. However, while processing samples from Lake St. Clair and Winnipeg we noticed increased formation of primer dimers generated mostly by original primer pair CYA359F-CYA781R. Based on our rigorous primer evaluation this primer pair does not contribute much to taxonomic recovery from environmental samples. Therefore, this primer pair can be excluded and replaced with modified primer cocktail CYA359Fd-CY781Rd-CYA781-euk.

Also, pooled libraries can be purified on E-gel SizeSelect II 2% agarose gels prior to sequencing to ensure complete primer dimer removal. Because PCR products amplified with M13-tailed primers are approaching the read length limit of Ion Torrent PGM with Hi-Q View chemistry, the fully automated workflow of Ion Torrent Chef/S5 instruments with Ext sequencing chemistry is a better fit for high-throughput analysis of environmental samples, as confirmed by our preliminary data.

We compared two approaches for identifying and quantifying phytoplankton assemblages. Clearly, both have limitations and biases; however, we argue that our NGS protocol offers a potentially new and powerful tool that can be better standardized than traditional microscopical analyses. We observed that microscopy results correlated with NGS data for Lake Erie samples collected in August 2015. However, taxonomic expertise is increasingly rare and costly, and as noted above, relies largely on morphological traits, many of which vary with environmental conditions and state of sample preservation [67,68]. Furthermore, few analysts remain current with evolving systematics and nomenclature, resulting in differences in classification and nomenclature. Not surprisingly, different labs vary in taxonomic expertise and the classification and counting methods used, and the few inter-lab comparisons published show series discrepancies amongst analysts; for example, there is a strong relationship between the number of microscope fields examined and the reported species diversity in a sample [69–72]. Unlike microscopy, NGS protocols and bioinformatics offer methods which can be easily standardized across different facilities, and identification results will rely on reference database, such as validated BOLD dataset DS-ECCSTAGE, containing sequences for most important bloom-forming cyanobacteria and eukaryotic algae found in Canadian waterbodies. While this method is therefore limited by the size of the reference database, this can be increased over time as new species are isolated and expanded to include other microorganisms such as bacteria, fungi, microzooplankton. Above all, the selection of the method(s) used clearly depends on the nature of the questions being addressed, however we believe that the best approach would ideally combine both NGS and microscopy, since the latter provides additional insight into field samples – inorganic and organic detritus, dead and dividing cells, colony configuration, cell size, presence of specialised cells etc. – not revealed by molecular analyses.

## Conclusions

Overall, this study resulted in the development of a method for the rapid assessment of environmental samples for major algal/cyanobacterial taxa present and, potentially harmful/invasive species. Curated BOLD reference dataset DS-ECCSTAGE containing 203 sequences representing 101 species can be used for identification of culture collection strains using Sanger sequencing or for NGS assessment of algal blooms or for routine monitoring of phytoplankton community diversity. We utilized this database along with mock community experiments to validate existing primer cocktails, and develop new NGS primer cocktail targeting 16S chloroplast rDNA in eukaryotic strains. Evaluation of commercial and in-house DNA extraction protocols indicated their general suitability for application to environmental samples. Simplified PCR setup produced comparable sequence data on environmental samples, while reducing input of DNA, which can be stored for additional analyses; Ion Torrent S5 and M13-tailed primer cocktails enable high-throughput processing of environmental samples. Developed methodology relies on publicly accessible BOLD 16S reference dataset DS-ECCSTAGE along with open source bioinformatic tools tailored to deal with taxa with low resolution. The HAB-ID toolset allows the reproducible detection of cyanobacteria and eukaryotic algae in environmental samples using high throughput and cost-effective NGS; moreover, its taxonomic scope can be further expanded with addition of more strains to the BOLD database, enabling its application for real-time monitoring of HABs temporal dynamics and species diversity issues in a wide range of waterbodies.

## Supporting information

**S1 Table. Strains used to assemble mock communities and microscopy results.**

**S2 Table. Environmental samples.**

**S3 Table. Annotated taxonomy file for OTUBlastParser.py.**

## Acknowledgements

This study was funded through Environment and Climate Change Canada under the Strategic Technology Applications of Genomics in the Environment (STAGE) program: “Rapid assessment of algal community composition and harmful blooms using DNA barcoding and Remote Sensing”. All sequencing analysis was done at the Canadian Centre for DNA Barcoding, Centre for Biodiversity Genomics, University of Guelph. We thank Evgeny Zakharov for advises on Tableau software; Janet Topan and Liuqiong Lu for assistance with Sanger sequencing; Bird’s laboratory at Université du Québec à Montréal (UQAM) and Kling’s laboratory at Algal Ecology & Taxonomy Inc. in Manitoba for microscopy analysis of algal cultures and environmental samples; Warwick Vincent for providing isolates from the Polar Cyanobacteria Culture Collection (U. Laval, Quebec, Canada).

## Author Contributions

**Conceptualization:** Susan B. Watson, Natalia V. Ivanova, L. Cynthia Watson

**Data curation:** L. Cynthia Watson, Natalia V. Ivanova

**Formal analysis:** Natalia V. Ivanova, L. Cynthia Watson

**Funding acquisition:** Susan B. Watson, Jérôme Comte, Natalia V. Ivanova, Timothy W. Davis, George S. Bullerjahn

**Investigation:** Natalia V. Ivanova, L. Cynthia Watson, Arusyak Abrahamyan, Susan B. Watson, Jérôme Comte

**Methodology:** Natalia V. Ivanova, Kyrylo Bessonov

**Software:** Kyrylo Bessonov

**Supervision:** Susan B. Watson, Jérôme Comte

**Validation:** Natalia V. Ivanova, L. Cynthia Watson, Susan B. Watson, Jérôme Comte

**Visualization:** Natalia V. Ivanova

**Writing – original draft:** Natalia V. Ivanova, L. Cynthia Watson, Jérôme Comte

**Writing – review & editing:** Natalia V. Ivanova, L. Cynthia Watson, Jérôme Comte, Kyrylo Bessonov, Arusyak Abrahamyan, Timothy W. Davis, George S. Bullerjahn, Susan B. Watson

